# Neural correlate of relief in the anterior cingulate cortex and ventral tegmental area: Neural correlate of relief in the ACC and VTA

**DOI:** 10.1101/102178

**Authors:** Thomas W. Elston, David K. Bilkey

## Abstract

Information gained during goal pursuit motivates adaptive behaviour. The anterior cingulate cortex (ACC) supports adaptive behaviour, but how ACC signals are translated into motivational signals remains unclear. Rats implanted in the ACC and ventral tegmental area (VTA), a dopaminergic brain area implicated in motivation, were trained to run laps around a rectangular track for a fixed reward, where each lap varied in physical effort (a 30cm climbable barrier). Partial directed coherence analysis of local field potentials revealed that ACC theta (4-12 Hz) activity increased as rats entered the barrier-containing region of the maze on trials when the barrier was absent, and predicted similar changes in VTA theta. This did not occur on effortful, barrier-present trials. These data suggest that ACC provides a top-down modulating signal which can influence the motivation with which to pursue a reward, and which may be, in our task, a neural correlate of relief.

## Introduction

Adaptation to changing circumstances is a common, critical consideration in nature (Darwin, 1865); however, the neural mechanisms enabling adaptive goal-directed behaviour have yet to be fully elucidated. The anterior cingulate cortex (ACC), a subregion of the anterior medial prefrontal cortex (AC; Vogt & Paxinos, 2014), has previously been shown to have a role in commitment to a course of action (Blanchard, Strait, & Hayden, 2015; Ma, Hyman, Phillips, & Seamans, 2014) and encoding value associated with physical effort (Cowen, Davis, & Nitz, 2012; Friedman et al., 2015; Hillman & Bilkey, 2010; Parvizi, Rangarajan, Shirer, Desai, & Greicius, 2013; Rudebeck, Walton, Smyth, Bannerman, & Rushworth, 2009; Walton, Devlin, & Rushworth, 2003). Furthermore, ACC signals received reward, predictions of future reward, and errors in such predictions (Bryden, Johnson, Tobias, Kashtelyan, & Roesch, 2011; Kennerly, Behrens, & Wallis, 2011). These signals are thought to facilitate the integration of new information into existing internal representations which can motivate and modify future behaviour (Bryden et al., 2011; Cavanagh, Frank, Klein, & Allen, 2010; Kolling, Behrens, Wittmann, & Rushworth, 2016). However, it remains unclear how ACC activity is translated into an adaptive, motivational signal.

The ACC and the anterior portions of the medial prefrontal cortex are potentially able to exert direct top-down control over the ventral tegmental area (VTA) via monosynaptic glutamatergic afferents (Beier et al., 2015; Carr & Sesack, 2000; Faget et al., 2016; Ferenczi et al., 2016; Gariano & Groves, 1988; Jo, Lee, & Mizumori, 2013; Taber, Das, & Fibiger, 1995). Moreover, in anaesthetised rats, ACC stimulation elicits burst firing in VTA dopamine neurons (Gariano & Groves, 1988). The VTA is a major source of dopamine to the basal ganglia and is hypothesized to convey a motivational signal indicating progress towards and value of a future goal (Hamid et al., 2016; Howe, Tierney, Sandberg, Phillips, & Graybiel, 2013; Morales & Margolis, 2017; Salamone & Correa, 2012). Large, transient bursts of single unit activity have also been reported in rodent in both ACC and VTA when an outcome is different than predicted (Bryden et al., 2011; Eshel, Tian, Bukwich, & Uchida, 2016). Strikingly, the magnitude of the activity increase correlates with the prediction-outcome error difference in both regions (Bryden et al., 2011; Eshel et al., 2015; Kennerly et al., 2011) suggesting that both these regions are involved in reward prediction error processes (Schultz, Dayan, & Montague, 1997). Thus, ACC top-down control over dopaminergic subcortical regions, such as the VTA, is a plausible mechanism by which ACC signals are translated into motivational signals.

Although no prior study has jointly examined the relationship between the ACC and VTA in the freely moving rat, prior investigations of anterior cortical (AC) and VTA communication suggest that these structures communicate via oscillations in the theta band (4-12 Hz). For example, firing in VTA dopamine neurons is modulated by the phase of AC theta oscillations and AC-VTA synchronization in the theta band is strongly correlated with working memory performance. However, to date, the precise relationship between the ACC and VTA and how these signals might shape future behaviour remains unknown.

To address this question directly, we monitored ACC and VTA local field potentials (LFPs) of rats trained to run laps around a rectangular track for a fixed reward, where each lap varied in physical effort. The effortful condition required rats to climb over a 30cm barrier while no barrier was present in the non-effortful condition. We examined task-dependent changes in theta (4-12 Hz) power and coherence and also assessed the directionality of communication between ACC and VTA via partial directed coherence (PDC), a multivariate autoregressive modelling approach which provides a frequency-resolved estimate of Granger causality (Baccalá & Sameshima, 2001; Boykin, Khargonekar, Carney, Ogle, & Talathi, 2012).

## Methods

### Subjects

Seven male Sprague Dawley rats (Hercus-Taieri Resource Unit) weighing 450-550 g were used in the study. Rats were single housed in translucent plastic cages containing pine chips and maintained on a 12 h light/dark cycle. All training and experimentation occurred during the light phase. After 2 weeks of daily handling and weighing, animals were food deprived of standard rat chow (Specialty Feeds) to no less than 85% of their free-feeding weight to promote interest in food reward during test phases. Water was available *ad libitum* in the home cage.

### Preoperative training

During the initial week of training, rats were individually habituated for 15 min/d to the experimental apparatus, a rectangular runway with a reward area (Figure 1A) which contained touchscreens, photobeams, and a retractable 30cm barrier which was mounted on a servo motor; the apparatus was controlled by a network of four Arduino (Arduino LLC, Somerville, MA, USA) microcontrollers. Cereal pellets (Coco Pop cereal; Kellogg's) were scattered throughout the maze to encourage exploration. In the second week of training, cereal pellets were only available at the reward site and rats were trained to run the maze in a unidirectional manner, starting at the bottom of the midstem (i.e. the startbox) and received a fixed reward (3 cocopops) in a distinct reward zone. Rats were prevented from reversing course in the maze by photobeam-controlled doors. The rats were not paused between trials and set their own pace, running in a continuous, uninterrupted manner.

**Figure 1.**
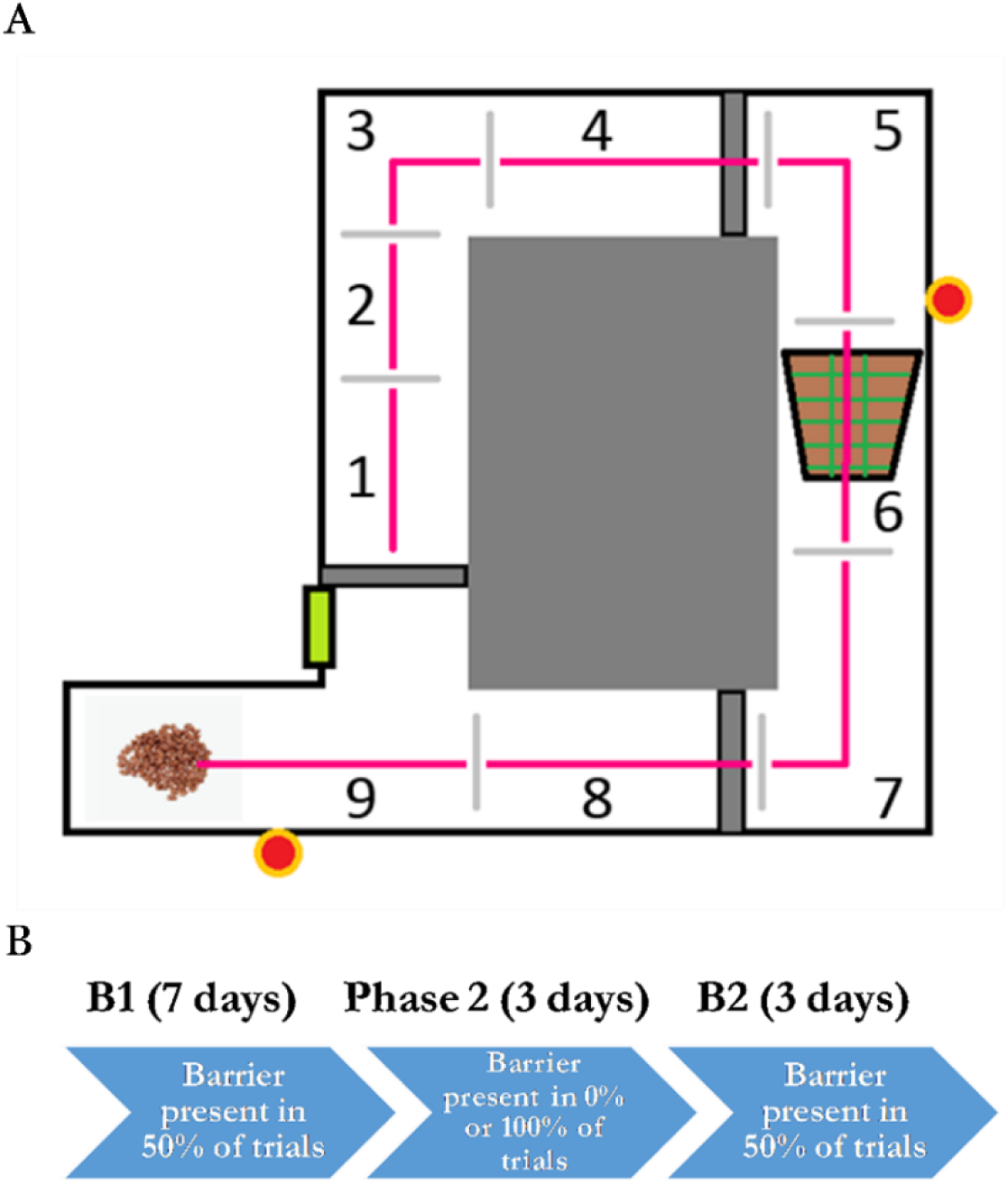
Schematic of maze and protocol. **A.** Rats initiated laps by pressing a wall-mounted touchscreen. After a 2-second delay, the startbox door opened and rats ran through regions 1-9. **B.** The experiment consisted of 3 phases: B1 (50% barrier), Phase 2 (either 0% barrier or 100% barrier), and B2 (50% barrier).

Once the rats were running unidirectionally around the apparatus, they were trained to press a wall-mounted touchscreen to release an adjacent starting gate and thereby initiate a new trial. This typically took two 15 minute training sessions or less. A two-second delay was then introduced between the initial press and the lowering of the start gate. Screen-press training was considered complete when rats ran >30 trials/session (2/min) across 3 consecutive training sessions. This typically took one week of training.

The final stage of training involved the introduction of a 30cm barrier into the apparatus which rats had to overcome on their route to the reward zone. The barrier was randomly inserted into the maze by a servo motor. Rats were cued as to whether the upcoming trial was an effortful, barrier trial or a non-effortful, flat trial by images delivered on the touch screen. The screen displayed three flashing green horizontal bars on a black background across the screen to indicate a barrier trial and a single non-flashing white bar on a magenta background to indicate a flat trial. To provide auditory camouflage for barrier insertion, the barrier was placed into the maze at the same instant the start gate was opened. Once rats were running >30 trials/session, sessions were extended such that each session required rats to run 120 laps composed of 60 trials without the barrier present (flat trials) and 60 barrier trials presented in pseudo-random order. Rats were considered fully trained when they were running 120 trials within 60 minutes (2/min).

### Surgery

Once animals were considered trained, five rats were anaesthetized under isoflurane and stereotaxically implanted in the ACC (AP: 2.7mm, ML: 0.4mm, DV: -1.8mm from dura) with 25 μm Formvar-coated nichrome wires (California Fine Wire) mounted on a 3D-printed adjustable microdrive assembly and the VTA (AP: -5.3mm, ML: 1.0mm, DV: - 8.2mm from dura) with a one non-moveable 127-μm-diameter, nickel-chromium coated wire. Two rats were also implanted at the same ACC coordinate with the microdrive assembly but with a second electrode (a fixed 127-μm-diameter, nickel-chromium coated wire) located in the dorsal CA1 subregion of the hippocampus (AP: -3.6mm, ML: 2.0mm, DV: -2.8mm from dura). The electrodes were grounded by soldering a wire to a jewellers screw implanted in the cerebellum. The assembly was fasted to the skull with jeweller’s screws and acrylic dental cement. Following the surgery, animals were allowed 10 days to recover during which they had *ad libitum* food and water. After 10 days, rats’ food was reduced to maintain the animal at ~85% of their free-feeding weight to optimize behaviour during the experiment.

### Postoperative training and protocol

After 10 days of recovery, rats were reintroduced to the maze with the head plugs connected to a tethered head stage that housed three light emitting diodes (LEDs) for tracking. Training resumed from the final preoperative stage; all 7 rats quickly recalled the task with no signs of postoperative motor impairment. Once each rat demonstrated three consecutive sessions of 120 trials in less than one hour, data acquisition began.

The data acquisition protocol consisted of a three stage, ‘ABA’ design (Figure 1B): seven days with protocol B1, during which the barrier was present in 50% of trials, three days Phase 2, during which touchscreen cues remained and the barrier was either 0% or 100% present, regardless of the cue, and three days of B2, in which the barrier was present in 50% of trials. Four rats experienced 0% barrier presence condition during Phase 2 and three rats experienced 100% barrier presence in Phase 2. All seven rats experienced B1 and B2 identically. It should be noted that the rats probably did not attend to the cues as there was no behavioural relevance to them and we found no behavioural or electrophysiological evidence that the cues themselves elicited any specific response.

### Electrophysiological recordings and data acquisition

Local field potentials were recorded using the dacqUSB multichannel recording system (Axona Ltd.). Local field potentials were low-pass filtered at 500 Hz, and sampled at 4800 Hz. The animals’ position was monitored by a ceiling-mounted video camera connected to a tracking system that monitored the LEDs mounted on the head stage. Tracking data was sampled at 50 Hz and made available to the dacqUSB system. Key events (e.g. trial initiation, the type of trial initiated, when rats got to the barrier region, and when rats reached the reward) were timestamped by inputs from a custom-built neural network of Arduino microcontrollers connected to a digital input-output port on the dacqUSB system.

### Data analysis

All analyses were conducted using MATLAB R2015b (The Mathworks, Boston, MA, USA). Power spectral density (PSD) data for the whole dataset were initially determined using using Welch’s estimate method, *pwelch* in MATLAB. Time resolved LFP power and coherence was then calculated via multi-taper spectrograms and coherograms (Mitra & Bokil, 2008) which used 3 tapers, a reading window of one second, and 85% overlap amongst windows. Data was linearized by dividing the maze path into 9 bins of 3-5cm in length and the position timestamps of rats entering and exiting a given region on a given lap on a given trial type were used to collect relevant pieces of the averaged 4-12 Hz segments of spectrograms and coheragrams which were then averaged for each maze region on each lap, separately. A similar approach was used to collect instantaneous speed data from each maze region for each lap on a given trial type. Task-related causal relationships between ACC-VTA and ACC-dCA1, linearly detrended LFPs were assessed with a partial directed coherence (PDC) algorithm, a frequency-resolved measure of Granger causality which uses multivariate autoregressive modelling to exploit the predictability of information in one brain area by past activity in another (Baccalá & Sameshima, 2001; Boykin, Khargonekar, Carney, Ogle, & Talathi, 2012).

Upon completion of the study, rats were deeply anaesthetized with isoflurane and recording sites were marked with direct current (2mA for 2 s) before transcardial perfusion. Electrode tracks and microlesions marking the electrode position were identified in 32µm thick sections of formalin-fixed tissue stained for Nissl substance (see Figure 2B).

**Figure 2.**
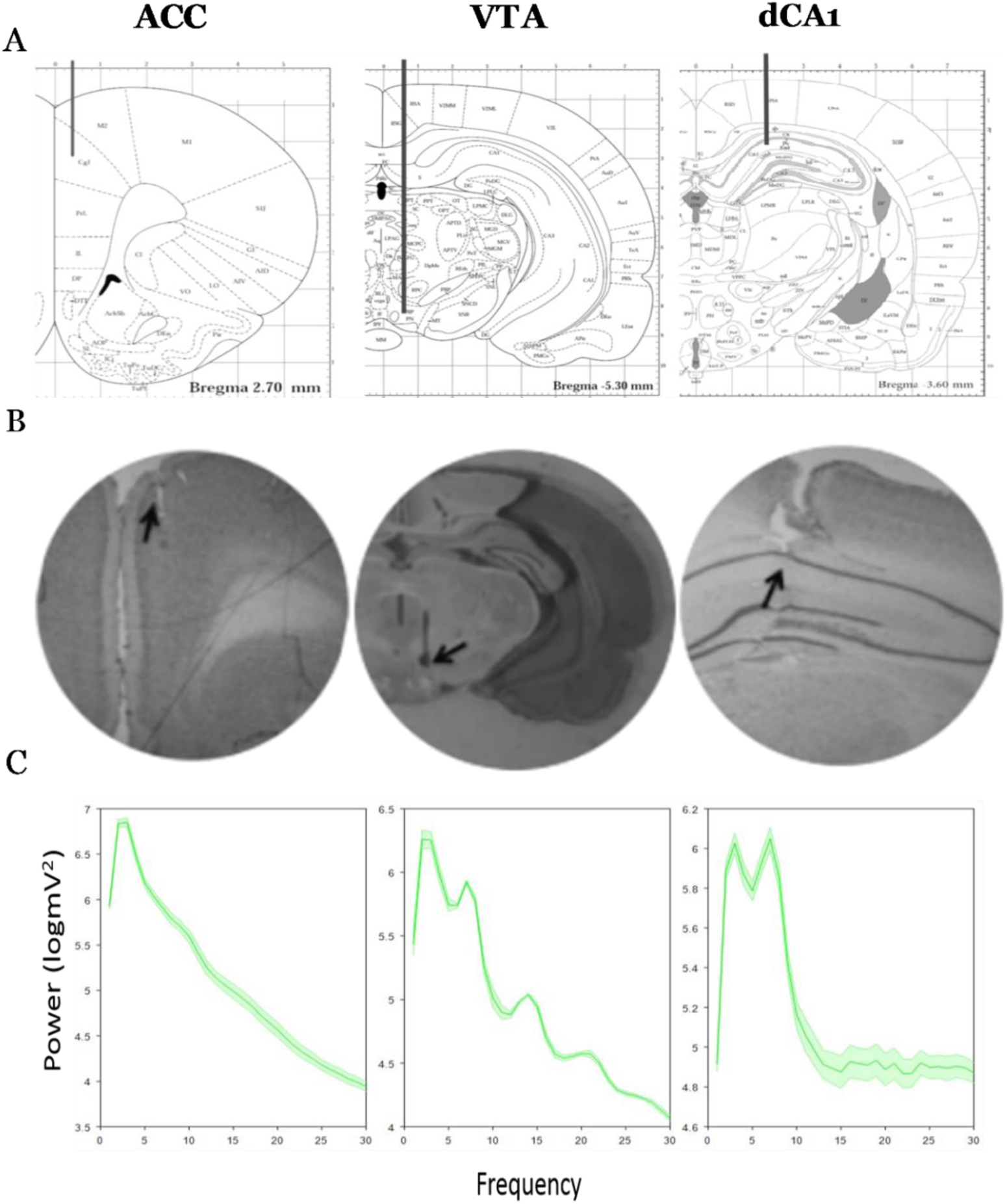
**A.** Plates from Paxinos & Watson (1997) indicating typical electrode placement in ACC (AP: 2.7mm, ML: 0.4mm, DV: -1.8mm from dura), VTA (AP: -5.3mm, ML: 1.0mm, DV: -8.2mm from dura), and dCA1 (AP: -3.6mm, ML: 2.0mm, DV: -2.8mm from dura). **B**. Histology indicating example electrode placement in ACC, VTA, and dCA1. **C.** Power spectrum densities (PSDs) determined from LFP via Welch’s method for ACC, VTA, and dCA1, in that order. The solid lines indicate the mean and the shaded areas are the standard errors.

## Results

### Task-modulated theta oscillations in ACC and VTA

Histology verified that electrodes were located in the regions of interest (Fig. 2 a-b). LFP patterns in ACC, VTA, and dCA1 were characteristically different with PSD functions from the ACC having a primary peak at low theta frequencies (3-5 Hz), while both VTA and dCA1 exhibited prominent peaks in both the high theta band (7-9 Hz) and low theta band (3-5 Hz; Figure 2C).

When the LFP data were examined on a region by region basis it was apparent that during phase B1, when the barrier was present 50% of the time, there was a marked difference in theta power, in both the VTA and ACC, and in the coherence between them, when the barrier-absent and barrier-present trials were compared (Figure 3). In barrier-absent trials there was a significant increase in low theta power and coherence that began once the animal approached the region of the maze immediately prior to where the barrier was usually present (region 5). Power increased through the barrier region and then was sustained above the level measured in the starting stem until the animal reached the reward zone (region 9). Power and coherence then remained high until a new trial was started. In barrier-present trials, however, theta power and coherence did not increase prior to or through the barrier region (p < .005 for regions 5-8, respectively, t-tests). Rather, levels remained supressed from the initial turn through until the reward region, at which point ACC and VTA theta power and coherence increased suddenly to become equivalent to that observed in barrier absent trials. This was a large effect (Hedge’s g > 1 for regions 5-8, respectively) and was observed across all animals, independently producing significant within-subject effects for every animal (regions 5-8, p < .0005, t-tests; see Figure 4 for all examples). No significant difference in ACC and VTA theta power or coherence was detected in the reward zone for any animal (region 9, all p > .05, t-tests).

**Figure 3.**
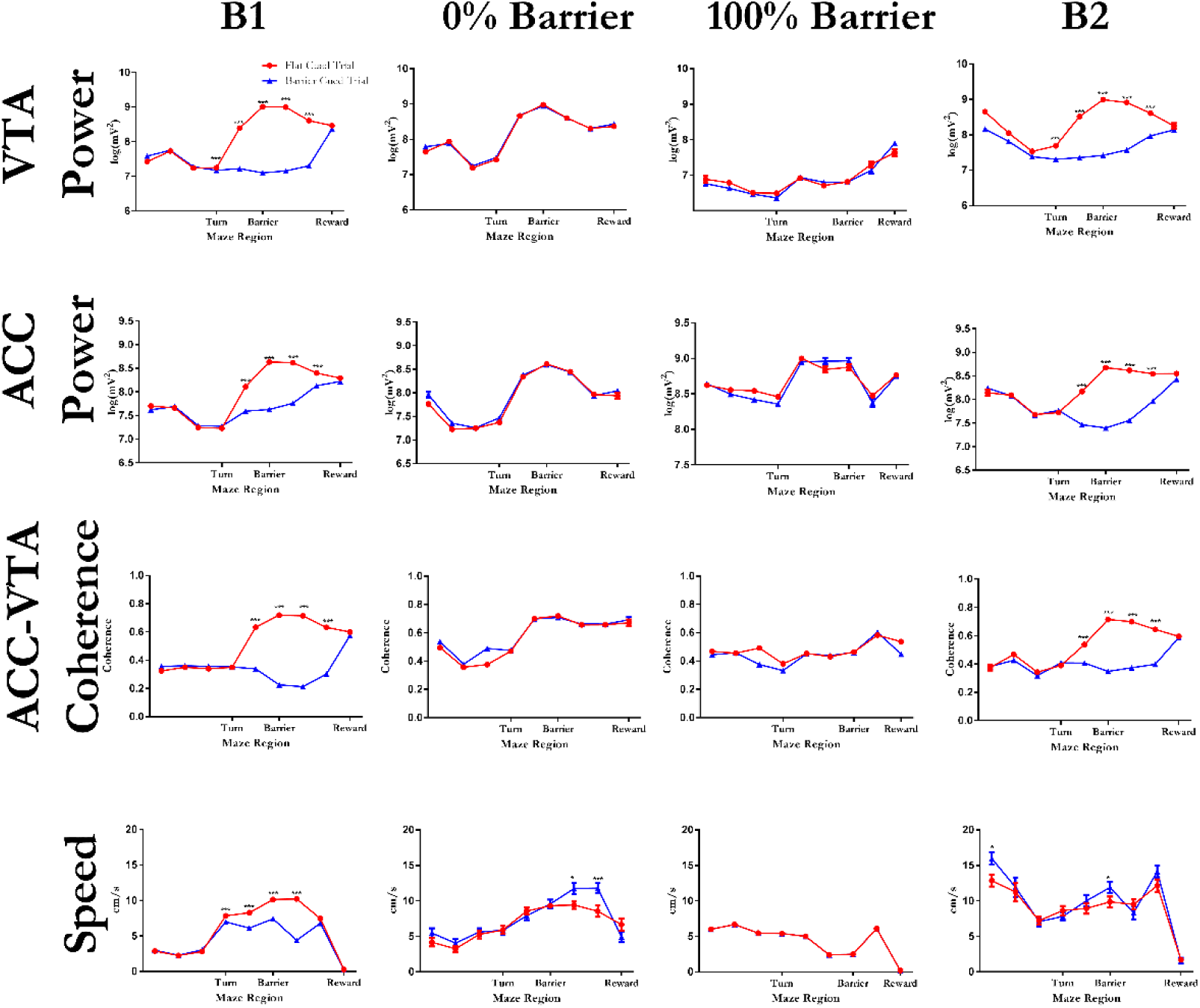
Enhanced theta (4-12Hz) power and coherence corresponds with the absence of the barrier. ACC and VTA time-resolved theta power, theta coherence, and running speed as a function of position for rat A (B1, 0%, and B2; mean ± SEM; n = 350 trials for B1, n = 150 trials for 0% barrier, and n = 150 trials for B2) and rat T (100% barrier; mean ± SEM; 150 trials for 100% barrier). The average power and coherence between 4 and 12 Hz for each region on each lap was used. *p < .05, **p < .005, ***p < .0005

**Figure 4.**
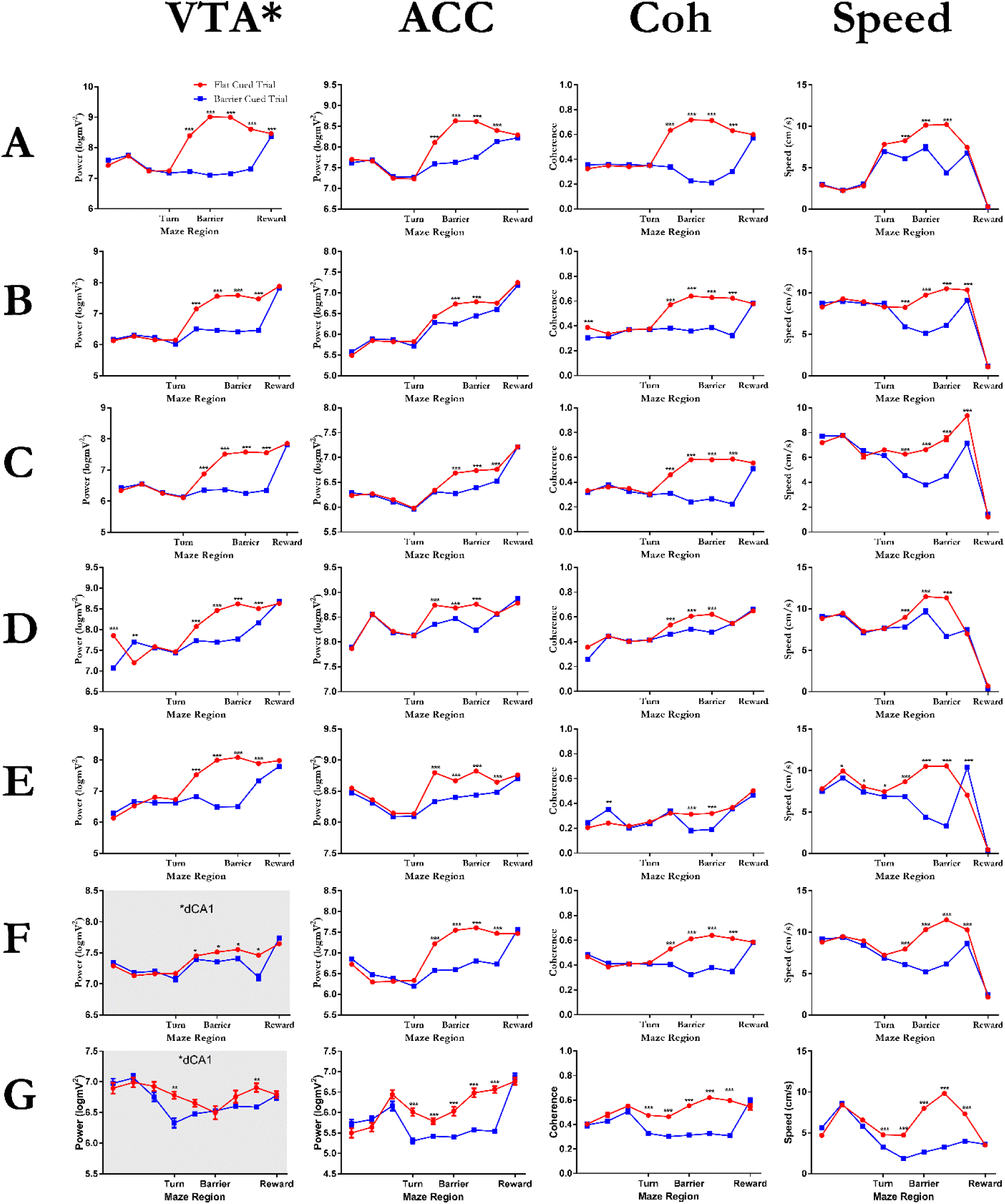
Enhanced theta (4-12Hz) power and coherence corresponds with the absence of the barrier in all rats. A. Data from all rats in phase B1. Rat name is listed on the left and brain region is listed along the top. *Data with grey background for rats F and G in the VTA column are dCA1.

In phase 2, the barrier was either not present for all trials (n = 4 animals) or was present for all trials (n=3 animals) irrespective of the cue on the touchscreen, which continued to randomly signal the two types of trials on a 50% basis. For animals where the barrier was absent, an abrupt increase in ACC and VTA theta power, similar to that which occurred in no-barrier trials in phase B1, was detected from regions 5 through 8 in all animals regardless of the trial type cued by the touch screen (Figure 5). For animals where the barrier was present on all trials, theta power remained low across these regions, replicating the effect of barrier-present trials in B1 (Figure 3). An analysis to determine whether the, now irrelevant and probably ignored, cue signal altered responses during phase 2 revealed no significant differences in ACC and VTA responses in regions 5 to 8 following the two cue types (all p > .05, t-tests).

**Figure 5.**
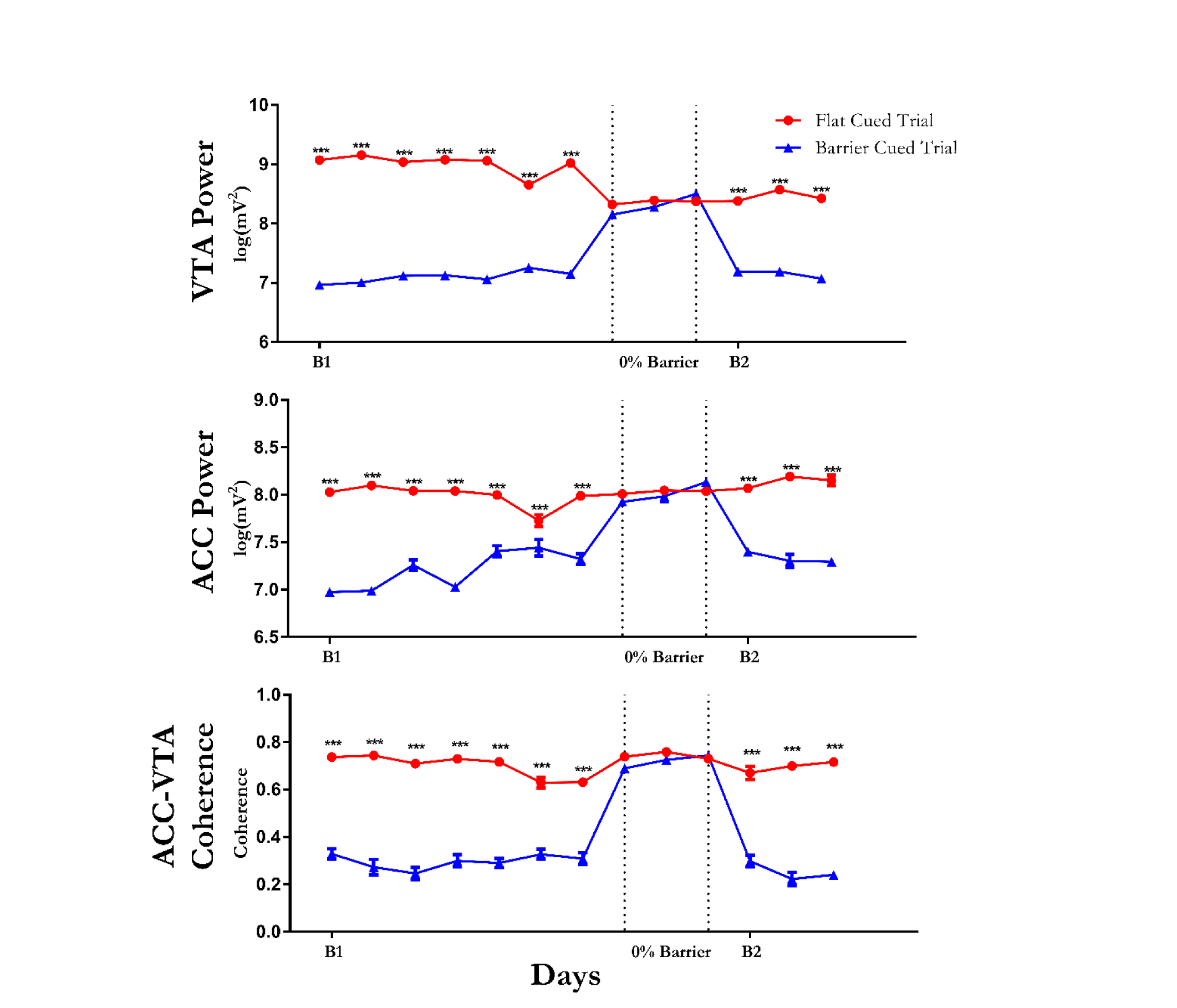
ACC and VTA theta and coherence responses to the presence or absence of the 30cm barrier in the barrier-containing region (region 6) of the maze were consistent across days and conditions. Data shown from rat A. VTA and ACC theta power and coherence were greater on trials when the barrier was present than when the barrier was absent across all days of B1 and B2 (p < .005, t-tests). *p < .05, **p < .005, ***p < .0005

In the third phase, B2, the barrier was again present on 50% of trials. In a result that replicated condition B1, an abrupt increase in running speed, ACC and VTA theta power, and coherence was detected in regions 5-8 in all animals when the barrier was absent (Figure 3; all p < .005, t-tests). On barrier trials, power and coherence remained suppressed until the reward region. Again, ACC and VTA theta power and coherence in the reward zone (region 9) did not depend on the trial type (p > .05, t-test). Analysis of responses across the multiple trials of each condition revealed that the significant modulation of ACC and VTA theta power and coherence by the presence or absence of the 30cm barrier in the barrier-containing region of the maze (region 6) was consistent across days and trial conditions (Figure 5).

### Task-modulated theta oscillations in ACC and dCA1

To assist in determining whether we were detecting a brain-wide phenomenon or activity that might be confined to specific regions, including VTA and ACC, we implanted two rats in both dCA1 and ACC and ran these animals through the procedure. As we had shown previously, in phase B1, we detected abrupt and sustained increases in ACC theta power from region 5 to 8 on no-barrier trials compared to barrier trials (p < .005, t-tests) in both animals (see Figures 4 and 6). In the second phase, when the barrier was absent on all trials, a similar abrupt increase in ACC theta power was detected regardless of the, likely ignored, cue (p > .05, t-tests). In phase B2, a significant difference in ACC theta power was detected in regions 5 to 8 when barrier and no-barrier trials were compared (all p < .005, t-tests) as we had observed previously. By comparison, recordings from dCA1 showed that there was a much smaller increase in theta power in regions 5-8 in B1 no-barrier trials. The difference between barrier and no-barrier trials was statistically significant (p < .05, t-test; Figure 4) in one animal but not the other (p > .05, t-test) in phase B1. In phase B2, however, no significant barrier-related changes were detected in dCA1 theta power in either animal (p > .05, t-tests, Figure 6). Despite the minimal effects seen in dCA1 theta power, ACC-dCA1 theta coherence was significantly modulated by the presence or absence of the 30cm barrier. In phases B1 and B2 dCA1-ACC theta coherence abruptly increased in region 5 and remained elevated through region 8 in no barrier trials compared to barrier trials (p < .005, t-tests). In phase 2, when the barrier was never present, a similar abrupt and sustained elevation in theta coherence, beginning in region 5, was detected regardless of the cued condition (p > .05, t-tests).

**Figure 6.**
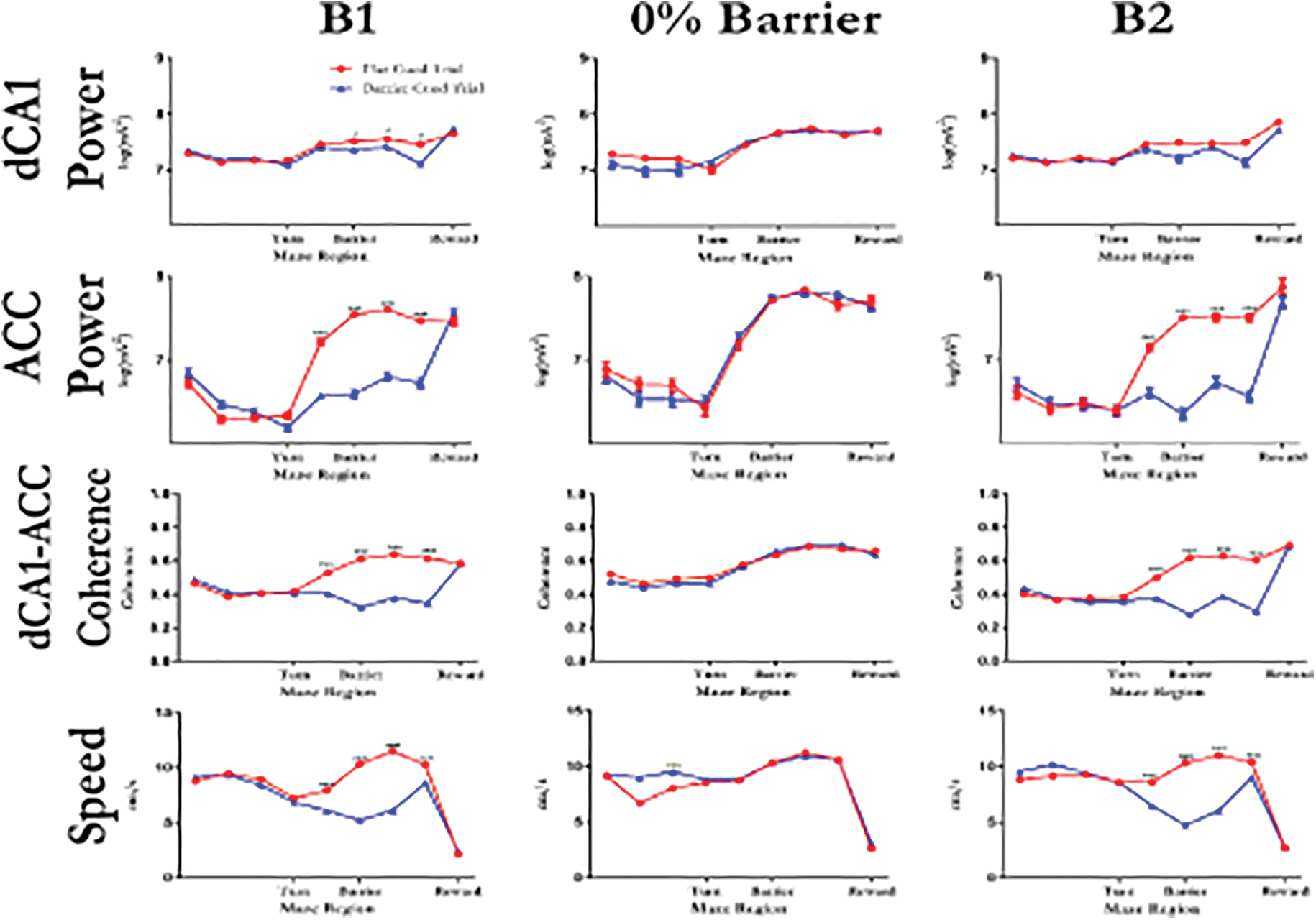
ACC but not dCA1 theta power was significantly modulated by the presence or absence of a 30cm barrier. Data are from Rat F. (mean ± SEM; n = 350 trials for B1, n = 150 trials for 0% barrier, and n = 150 trials for B2) *p < .05, **p < .005, ***p < .0005

### Relationship to running speed

We were concerned that our results may have simply been an artefact of changes in running speed; indeed, at first glance, the patterns of power and coherence seemed to parallel changes in speed (Figure 3). However, upon closer inspection, it’s clear that running speed and LFP power and coherence are not always changing together. For instance, an increase in running speed occurred in region 4 before the increase in power and coherence was observed (region 5). Furthermore, while running speed is near zero in the reward region, LFP power and coherence is elevated. To formally assess the relationship between running speed and LFP dynamics, we initially conducted correlations (Pearson’s R) between the instantaneous running speed and instantaneous theta power (4-12Hz) in ACC, VTA, and dCA1, respectively, across all sessions and all regions of the maze separately for each rat, considering only instances when rats were moving (e.g. speed > 1). All correlation coefficients were near zero and none were significant (all r < 10-2, all p > .05; see Figure 7A-C and Table 1), suggesting that speed and LFP power were modulated independently. To further examine the relationship between running speed and LFP power under different effort conditions, we conducted analyses of covariance (ANCOVA) with instantaneous ACC, VTA, or dCA1 theta power in the barrier region as the dependent variable, with the corresponding instantaneous running speed as a covariate, and the presence or absence of the barrier as the independent variable for all sessions of phase B1 for each animal, separately. Resultant ANCOVAs revealed a significant relationship between VTA, ACC, and dCA1 theta power in the barrier region and running speed in the barrier region (see Table 2). This differs from lack of correlations observed across the full region as described above. After statistically controlling for running speed as a covariate, the presence or absence of the barrier remained a highly significant factor in the ACC and VTA, but not dCA1, LFP power in all rats (all p < .0005; see Table 3). These results indicate that instantaneous running speed and instantaneous VTA and ACC theta power are independently modulated by the effort condition, whereas dCA1 theta power was dependent on running speed.

**Figure 7.**
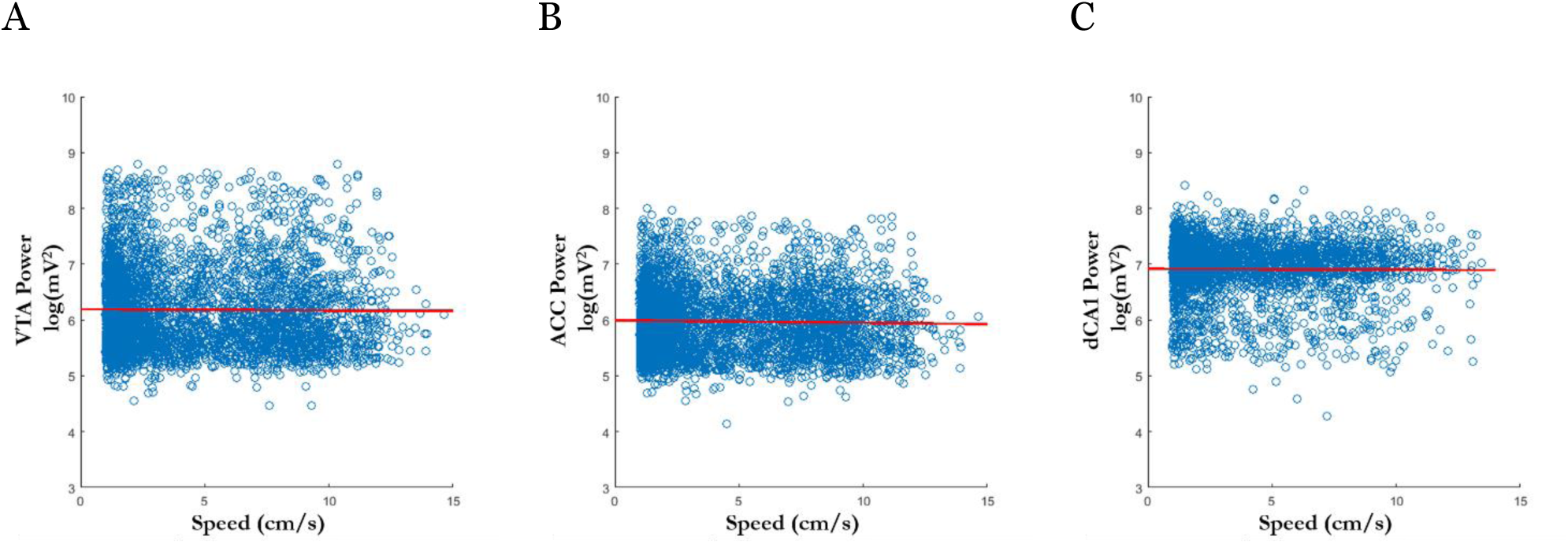
Relationships between running instantaneous speed and instantaneous LFP theta power while rats moved > 1 cm/s. **A.** Data from rat A showing the correlations between running speed and VTA theta power. **B.** Data from rat A showing the correlations between running speed and ACC theta power. **C.** Data from rat F showing the correlations between running speed and dCA1 theta power, respectively.

**Table 1.** Running speed – LFP theta (4-12 Hz) power correlation statistics for all rats.

| Table 1.<br>Correlations Between Running Speed and LFP Theta (4-12 Hz) Power by Brain Region and Rat |  |  |  |
| --- | --- | --- | --- |
| Rat | VTA and Running Speed | ACC and Running Speed | dCA1 and Running Speed |
| A | $r(16693) = -1.76 \times 10^{-5}$, $p > .05$ | $r(16693) = -1.90 \times 10^{-3}$, $p > .05$ | |
| B | $r(96303) = -1.70 \times 10^{-4}$, $p > .05$ | $r(96303) = -6.49 \times 10^{-4}$, $p > .05$ | |
| C | $r(106428) = -1.53 \times 10^{-4}$, $p > .05$ | $r(106428) = -6.09 \times 10^{-4}$, $p > .05$ | |
| D | $r(145577) = 2.40 \times 10^{-3}$, $p > .05$ | $r(145577) = -1.80 \times 10^{-4}$, $p > .05$ | |
| E | $r(159581) = 4.00 \times 10^{-3}$, $p > .05$ | $r(159581) = 6.4 \times 10^{-3}$, $p > .05$ | |
| F | | $r(78535) = 3.55 \times 10^{-4}$, $p > .05$ | $r(78535) = 2.90 \times 10^{-4}$, $p > .05$ |
| G | | $r(123664) = 5.10 \times 10^{-3}$, $p > .05$ | $r(123664) = 5.8 \times 10^{-3}$, $p > .05$ |

**Table 2.** ANCOVA results indicating significant covariance between theta power and running speed in the barrier region.

| Table 2. |  |  |  |
| --- | --- | --- | --- |
| B1 ANCOVA Results for Theta (4-12 Hz) Power and Speed Relationship by Brain Region and Rat |  |  |  |
| Rat | VTA Theta Power | ACC Theta Power | dCA1 Theta Power |
| A | $F_{1,116828} = 6.95 \times 10^3, p < 1.00 \times 10^{-200}$ | $F_{1,82363} = 3.52 \times 10^3, p < 1.00 \times 10^{-200}$ | |
| B | $F_{1,35669} = 4.59 \times 10^3, p < 1.00 \times 10^{-200}$ | $F_{1,35669} = 2.30 \times 10^3, p < 1.00 \times 10^{-200}$ | |
| C | $F_{1,52251} = 2.65 \times 10^3, p < 1.00 \times 10^{-200}$ | $F_{1,52251} = 6.83 \times 10^2, p = 1.25 \times 10^{-149}$ | |
| D | $F_{1,40881} = 3.81 \times 10^3, p < 1.00 \times 10^{-200}$ | $F_{1,40881} = 7.51 \times 10^3, p < 1.00 \times 10^{-200}$ | |
| E | $F_{1,73924} = 9.53 \times 10^3, p < 1.00 \times 10^{-200}$ | $F_{1,73924} = 2.91 \times 10^3, p < 1.00 \times 10^{-200}$ | |
| F | | $F_{1,45583} = 2.01 \times 10^3, p < 1.00 \times 10^{-200}$ | $F_{1,82363} = 179.25, p = 8.41 \times 10^{-41}$ |
| G | | $F_{1,82363} = 104.43, p < 1.00 \times 10^{-200}$ | $F_{1,82363} = 4.96 \times 10^3, p < 1.00 \times 10^{-200}$ |

**Table 3.** ANCOVA results for factor barrier presence as a significant predictor of theta power in ACC and VTA in the barrier region after adjusting for running speed as a covariate.

| Table 3.<br>B1 ANCOVA Results for Barrier Presence After Controlling for Run Speed by Brain Region and Rat |  |  |  |
| --- | --- | --- | --- |
| Rat | VTA Theta Power | ACC Theta Power | dCA1 Theta Power |
| A | $F_{1,116828} = 9.17, p = 2.50 \times 10^{-3}$ | $F_{1,82363} = 4.38, p = 5.00 \times 10^{-3}$ | |
| B | $F_{1,35669} = 1.77 \times 10^3, p < 1.00 \times 10^{-200}$ | $F_{1,35669} = 493.64, p = 1.26 \times 10^{-108}$ | |
| C | $F_{1,52251} = 2.52 \times 10^3, p < 1.00 \times 10^{-200}$ | $F_{1,52251} = 5.79 \times 10^3, p < 1.00 \times 10^{-200}$ | |
| D | $F_{1,40881} = 2.19 \times 10^3, p < 1.00 \times 10^{-200}$ | $F_{1,40881} = 507.99, p = 8.38 \times 10^{-112}$ | |
| E | $F_{1,73924} = 2.05 \times 10^3, p < 1.00 \times 10^{-200}$ | $F_{1,73924} = 169.73, p = 9.35 \times 10^{-39}$ | |
| F | | $F_{1,45583} = 661.15, p = 9.11 \times 10^{-145}$ | $F_{1,82363} = 3.43, p > .05$ |
| G | | $F_{1,82363} = 7.23, p = 3.26 \times 10^{-4}$ | $F_{1,82363} = 1.37, p > .05$ |

### ACC top-down influence over dCA1 and VTA is modulated by barrier presence

We then used a partial directed coherence (Baccalá & Sameshima, 2001) procedure following Boykin, Khargonekar, Carney, Ogle, & Talathi (2012)’s method to determine if there was evidence of signal directionality in linearly detrended LFPs recorded simultaneously from ACC and VTA and ACC and dCA1 as the animal passed through the barrier-containing region (region 6) of the maze. The mean partial directed coherence (PDC) was calculated in a trial-by-trial manner for each condition for data in the theta range (4-12 Hz). An ANOVA examining the resultant PDC between ACC and VTA in B1 revealed significant main effects of directionality (F_1,1194_ = 719.45, p = 2.01 × 10^-124^) and barrier condition (F_1,1194_ = 121.57, p = 5.46 × 10^-27^) as well as a significant directionality × barrier interaction effect on ACC-VTA theta PDC responses (F_1,1194_ = 151.02, p = 9.03 × 10^-33^; see Figure 8A; see Table 4 for data from all rats). An ANOVA considering B2 returned nearly identical results (see Table 5). This interaction is a result of ACC → VTA PDC being higher when the barrier was absent and decreasing in barrier trials while VTA→ACC PDC remained low and unchanging across conditions. Similarly, an ANOVA examining PDC between ACC and dCA1 across all trials and conditions revealed significant main effects of directionality (F_1,1797_ = 207.32, p = 5.3 × 10^-44^) and barrier presence (F_1,1797_ = 233.09, p = 1.43 × 10^-49^) as well as a significant directionality × barrier interaction effect on ACC-dCA1 theta PDC responses (F_1,1797_ = 35.69, p = 2.78 × 10^-9^; see Figure 8B; Tables 4 and 5). This interaction was a result of dCA1→ACC PDC decreasing to a greater degree when the barrier was present compared to ACC→dCA1 PDC. To verify the integrity of our PDC modelling, we scrambled the recorded signals from both ACC and VTA and conducted the trial-by-trial PDC analysis again. In every case the mean of the permutated PDC distribution was significantly weaker than the non-permuted data (all PDC < 3×10^-5^, all p < 3.76×10^-36^), losing evidence for directionality.

**Table 4.** Individual rat barrier-region PDC ANOVA main and interaction effects by implant type in condition B_1_.

| Table 4.<br>Barrier Region PDC B1 ANOVA Results by Rat, Implant Type, and Effect |  |  |  |  |
| --- | --- | --- | --- | --- |
| Rat | Implant | Directionality | Barrier Presence | DxBP Interaction |
| A | ACC-VTA | $F_{1,1194} = 362.56, p = 5.57 \times 10^{-72}$ | $F_{1,1194} = 33.91, p = 7.71 \times 10^{-9}$ | $F_{1,1194} = 46.82, p = 1.16 \times 10^{-11}$ |
| B | ACC-VTA | $F_{1,1194} = 719.45, p = 2.01 \times 10^{-124}$ | $F_{1,1194} = 121.57, p = 5.46 \times 10^{-27}$ | $F_{1,1194} = 151.02, p = 9.03 \times 10^{-33}$ |
| C | ACC-VTA | $F_{1,1194} = 351.65, p = 4.24 \times 10^{-74}$ | $F_{1,1194} = 23.56, p = 6.78 \times 10^{-34}$ | $F_{1,1194} = 54.22, p = 3.96 \times 10^{-29}$ |
| D | ACC-VTA | $F_{1,1194} = 402.95, p = 5.32 \times 10^{-75}$ | $F_{1,1194} = 14.08, p = 2.00 \times 10^{-4}$ | $F_{1,1194} = 341.11, p = 1.00 \times 10^{-4}$ |
| E | ACC-VTA | $F_{1,1194} = 352.77, p = 2.78 \times 10^{-68}$ | $F_{1,1194} = 20.55, p = 3.42 \times 10^{-7}$ | $F_{1,1194} = 88.45, p = 4.64 \times 10^{-22}$ |
| F | ACC-dCA1 | $F_{1,1194} = 242.50, p = 1.44 \times 10^{-50}$ | $F_{1,1194} = 310.47, p = 6.67 \times 10^{-63}$ | $F_{1,1194} = 30.40, p = 5.02 \times 10^{-8}$ |
| G | ACC-dCA1 | $F_{1,1194} = 528.33, p = 2.15 \times 10^{-99}$ | $F_{1,1194} = 273.99, p = 2.53 \times 10^{-56}$ | $F_{1,1194} = 70.1, p = 1.36 \times 10^{-16}$ |

**Table 5.** Individual rat barrier-region PDC ANOVA main and interaction effects by implant type in condition B_2_.

| Table 5.<br>Barrier Region PDC B2 ANOVA Results by Rat, Implant Type, and Effect |  |  |  |  |
| --- | --- | --- | --- | --- |
| Rat | Implant | Directionality | Barrier Presence | DxBP Interaction |
| A | ACC-VTA | $F_{1,596} = 118.17, p = 1.10 \times 10^{-10}$ | $F_{1,596} = 12.88, p = 4.00 \times 10^{-4}$ | $F_{1,596} = 12.47, p = 4.00 \times 10^{-4}$ |
| B | ACC-VTA | $F_{1,596} = 628.74, p = 2.78 \times 10^{-95}$ | $F_{1,596} = 129.12, p = 3.21 \times 10^{-27}$ | $F_{1,596} = 175.93, p = 2.28 \times 10^{-35}$ |
| C | ACC-VTA | $F_{1,596} = 482.87, p = 7.69 \times 10^{-79}$ | $F_{1,596} = 125.47, p = 1.46 \times 10^{-26}$ | $F_{1,596} = 92.69, p = 1.74 \times 10^{-20}$ |
| D | ACC-VTA | $F_{1,596} = 492.48, p = 1.00 \times 10^{-82}$ | $F_{1,596} = 12.69, p = 4.00 \times 10^{-4}$ | $F_{1,596} = 122.42, p = 1.37 \times 10^{-25}$ |
| E | ACC-VTA | $F_{1,596} = 416.22, p = 6.88 \times 10^{-70}$ | $F_{1,596} = 50.60, p = 7.80 \times 10^{-15}$ | $F_{1,596} = 8.95, p = 2.90 \times 10^{-3}$ |
| F | ACC-dCA1 | $F_{1,596} = 197.36, p = 7.36 \times 10^{-39}$ | $F_{1,596} = 100.61, p = 5.66 \times 10^{-22}$ | $F_{1,596} = 17.18, p = 3.89 \times 10^{-5}$ |
| G | ACC-dCA1 | $F_{1,596} = 245.74, p = 1.26 \times 10^{-46}$ | $F_{1,596} = 233.99, p = 8.45 \times 10^{-45}$ | $F_{1,596} = 88.32, p = 1.18 \times 10^{-19}$ |

**Figure 8.**
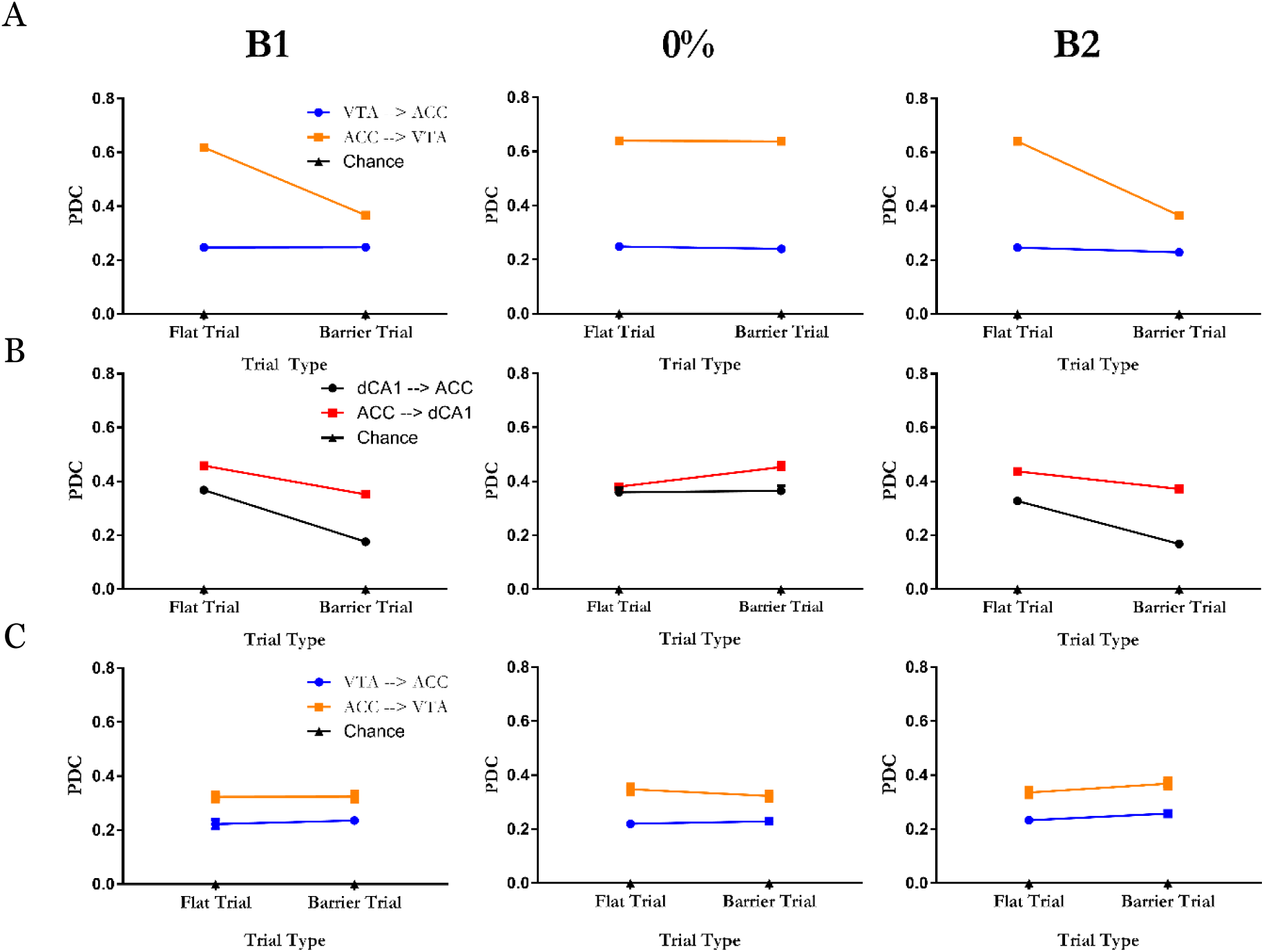
ACC-VTA and ACC-dCA1 partial directed coherence (PDC) in the barrier region. Data indicates mean ± SEM. Some error bars were too small for display. **A.** ACC → VTA and VTA → ACC theta PDC in the barrier region across the three phases. **B.** ACC→ dCA1 and dCA1 → ACC theta PDC in the barrier region across the three phases. **C.** ACC→VTA and VTA‐->ACC PDC in the initial region (region 2) across the three phases. and C are data from rat B; B are from rat G.

To determine if this PDC effect was specific to the barrier region, we also assessed the relationship between the ACC and VTA and ACC and dCA1 as animals passed through the initial region of the apparatus (region 2). An ANOVA examining the resultant PDC between ACC and VTA in this region revealed a significant main effect of directionality (F_1,1194_ = 286.65, p < × 10^-200^) but there was no significant effect of barrier condition (F_1,1194_ = 3.46, p > .05) and no directionality × barrier interaction (F_1,1194_ = 0.76, p > .05; see Figure 8C; see Table 6 for data from all rats). Similar results were obtained for the ACC-dCA1 PDC ANOVAs (see Table 6), These data indicate that in the initial portion of the apparatus, where no variation in task-demands ever occurred, ACC leads VTA and dCA1 activity, respectively, but without the large increase that occurs when the animal discovers the barrier is absent from the barrier-containing region of the apparatus.

**Table 6.** Individual rat initial segment PDC ANOVA main and interaction effects by implant type in condition B_2_.

| Table 6.<br>Initial Segment PDC B1 ANOVA Results by Rat, Implant Type, and Effect |  |  |  |  |
| --- | --- | --- | --- | --- |
| Rat | Implant | Directionality | Barrier Presence | DxBP Interaction |
| A | ACC-VTA | $F_{1,1194} = 4.65, p = .032$ | $F_{1,1194} = .1117, p > .05$ | $F_{1,1194} = 2.959, p > .05$ |
| B | ACC-VTA | $F_{1,1194} = 99.43, p = 9.52 \times 10^{-22}$ | $F_{1,1194} = .4118, p > .05$ | $F_{1,1194} = 1.264, p > .05$ |
| C | ACC-VTA | $F_{1,1194} = 616.59, p = 4.13 \times 10^{-110}$ | $F_{1,1194} = 1.109, p > .05$ | $F_{1,1194} = 3.642, p > .05$ |
| D | ACC-VTA | $F_{1,1194} = 292.23, p = 8.75 \times 10^{-59}$ | $F_{1,1194} = .2456, p > .05$ | $F_{1,1194} = 2.595, p > .05$ |
| E | ACC-VTA | $F_{1,1194} = 1311.0, p = 7.30 \times 10^{-194}$ | $F_{1,1194} = 3.483, p > .05$ | $F_{1,1194} = 2.667, p > .05$ |
| F | ACC-dCA1 | $F_{1,1194} = 306.85, p = 2.48 \times 10^{-61}$ | $F_{1,1194} = .0912, p > .05$ | $F_{1,1194} = 2.823, p > .05$ |
| G | ACC-dCA1 | $F_{1,1194} = 266.10, p = 4.06 \times 10^{-54}$ | $F_{1,1194} = 3.264, p > .05$ | $F_{1,1194} = 3.636, p > .05$ |

## Discussion

We investigated the influence of physical effort on VTA, ACC, and dCA1 theta (4-12 Hz) power, coherence, partial directed coherence (PDC), and running speed as rats ran a rectangular track requiring varying levels of effort to reach a fixed reward. PDC analysis of task-related casual relationships between brain areas revealed significant ACC → VTA and ACC → dCA1 directionality, the magnitude of which was modulated by the presence or absence of the barrier. These PDC results show that ACC activity is predictive of VTA and dCA1 responses and are consistent with prior reports showing cortical influence over VTA and dCA1. For example, Beier et al. (2015) found that the VTA dopaminergic neurons projecting to NAcc are directly regulated by anterior-cortical-VTA glutamatergic afferents and that rodents will self-stimulate those projections. Additionally, ACC control over the VTA is strongly implicated in inhibition of cocaine seeking (Navailles, Guillem, Vouillac-Mendoza, & Ahmed, 2015). Moreover, Rajasethupathy et al. (2015) found that ACC projections to dCA1 mediate memory retrieval and Ito et al. (2015) implicated ACC influence over dCA1 in directing goal-oriented place-cell activity. One interpretation of these findings is that ACC top-down projections to the VTA and dCA1 (Ferenczi et al., 2016; Fujisawa & Buzsaki, 2011; Taber et al., 1995) underlie a circuit via which cortical representations can update spatial and motivational brain states to guide future behaviour.

Additionally, we found that ACC and VTA, but not dCA1, theta power, ACC-VTA coherence, and, surprisingly, ACC-dCA1 coherence increased as the rat entered the barrier-containing region of the maze on trials when the barrier was absent. In contrast, theta signals did not increase to the same degree across the same region of space when the barrier was present. Once the animal reached the reward region, however, ACC and VTA theta power and coherence as well as ACC-dCA1 theta coherence were independent of the effort condition.

Rats’ running speeds also increased significantly in the barrier region when the barrier was absent. This may have been due to the reduced physical demands of the task or, alternatively, may have been driven by increased motivation signals, perhaps provided in a top-down manner from the ACC. In either case, one consequence of this relationship is that changes in running speed are potentially confounded with some of the electrophysiological effects that we observed. Our analyses indicate, however, that the changes in electrophysiology we observed are over and above that which would be predicted from the changes in running speed *per se*. First, correlations between instantaneous power and instantaneous running speed when compared across all regions, were minimal and non-significant, suggesting that speed and power were independently driven by the effort condition. Second, and most importantly, after statistically controlling for speed as a covariate in an ANCOVA, the effort condition remained a highly significant factor in ACC and VTA (but not dCA1). Third, if the theta power changes were movement artefacts, it is difficult to explain why activity in the barrier region during the barrier condition, where rats would vigorously leap atop the barrier, was generally equal to that recorded in the initial stem of the apparatus where there was never a barrier. Therefore, rather than being a secondary consequence of behaviour change, our data appear to represent fundamental differences in underlying neural activity that are responsive to the effort condition.

Our data may reflect the operation of several different mechanisms, with the possibility that more than one of these operates simultaneously. Two of these include prediction error (Pearce & Hall, 1980) and negative reinforcement (Skinner, 1938). Prediction error (PE) theory posits that, based on prior experience and outcomes, the brain predicts what will happen in a similar situation and elicits a strong signal when an outcome violates that prediction. In our study, theta power in the ACC and VTA increased markedly when the barrier was absent. Framed as a prediction error effect, this would suggest that the baseline prediction might have been the presence of the barrier, such that the absence of the barrier was ‘unexpected’. This corresponds to the notion of a positively signed prediction error (Eshel et al., 2015; Kennerly et al., 2011), but is inconsistent with other reports of unsigned, ‘surprise’, prediction errors (Bryden et al., 2011; Hayden, Heilbronner, Pearson, & Platt, 2011). Were these signals true prediction error signals, the presence or absence of the barrier in the B1 condition, when the barrier was present on 50% of trials, should be equally surprising with each trial type eliciting a transient response.

Negative reinforcement (NR) models suggest that the removal of an aversive stimulus, such as a barrier, is itself a reinforcer (Skinner, 1938). Typical NR studies require subjects to emit an instrumental response to remove or avoid an aversive stimulus (Qi et al., 2016) and adapt future behaviour (Oleson, Gentry, Chioma, & Cheer, 2012). Although our study is slightly different from a typical NR design in that the omission of the barrier did not require an instrumental response, the increase in theta power and coherence along with running speed is consistent with the core notion of NR: the absence of a barrier, when it could have been present, is reinforcing. Both of these interpretations would predict that, in the phase 2, barrier-absent condition, the magnitude of the power and coherence signal would decrease in the barrier region over repeated trials as the animals’ predictions were modified to accept the new barrier probability (0%). The data presented in figure 5 suggest that this was not the case, at least within the time window tested. Thus, future experiments are required to resolve whether the time course over which these neural mechanisms respond to environmental change corresponds to that observed in behavioural change, such as extinction.

A related interpretation is suggested by human fMRI experiments. These dissociated the ACC’s response to surprising, presumably PE-inducing events, from its response to information indicating that the organism’s underlying predictive model must be updated (Kolling et al., 2016; O’Reilly et al., 2013). According to this model, the responses of the ACC and the structures it influences, such as the VTA, and, ultimately, behavioural changes are greatest when task-requirements are different than expected, such as a barrier being absent when it has previously been present. This is different from PE in that PEs are thought to signal one-off, ‘surprising’ events whereas Kolling et al. (2016) and O’Reilly et al. (2013) suggest these ACC signals are updating the more fundamental processing framework underlying PEs. This interpretation reconciles the key aspects of prediction error and negative reinforcement in that VTA and ACC theta power as well as running speed abruptly increased when the rat finds that the barrier is absent, when it has been present in the past. Future experiments are required to verify an assumption of this explanation which, similar to a PE account, assumes the animals’ default task model includes the presence of the barrier.

It is also possible that our findings can also be explained as a neural correlate of ‘relief’, in that the avoidance of an aversive outcome, such as climbing a barrier, is relieving. Such a description of relief is consistent with human psychological conceptions of relief (Carver, 2009; Deutsch, Smith, Kordts-Freudinger, & Reichardt, 2015; Sweeny & Vohs, 2012). Since relief is thought to occur only when circumstances or outcomes are better than they might have been, the concept of relief synthesises the key aspects of negative reinforcement, positive prediction error, and internal task-state monitoring-and-updating without many of the caveats of other interpretations. Thus, characterisation of our findings as a neural correlate of relief is perhaps a more parsimonious description of a process by which internal task representations are altered, specifically when an aversive task-requirement, such as climbing a barrier, is avoided.

Our data integrates and extends the evidence indicating a close link between cortical states and activity in dopaminergic brain areas (Beier et al., 2015; Fujisawa & Buzsaki, 2011; Taber et al., 1995; Wu et al., 2013) and complements demonstrations of cortical control of other affective states, such as fear (Karalis et al., 2016; Likhtik, Stujenske, Topiwala, Harris, & Gordon, 2014), apathy (Moretti & Signori, 2016; Onoda & Yamaguchi, 2015), and pain (Navratilova & Porreca, 2014; Vogt, 2005). The idea of a continuously sampling ACC which tracks changes in task requirements complements reports of a similarly adaptive mesolimbic dopamine signal (Hamid et al., 2016; Howe et al., 2013) and provides an indication of at least some of the pathways via which this latter response could be modulated. Together, these data suggest that cortical and dopaminergic states modulate the motivation to pursue a reward in response to changing task requirements, enabling organisms to behave flexibly in order to optimize available opportunities in uncertain, changing environments (Hayden, Pearson, & Platt, 2011). Integrating the theories of prediction error, negative reinforcement, and task-requirement monitoring, we conclude that our findings may reflect a neural correlate of relief: the avoidance of a barrier, when it has been present in the past, is relieving. Future work will expand the mechanistic insight of this phenomenon by examining how changes in the ACC to VTA projection map onto changes in behavioral outcomes.

## Acknowledgements

The Marsden Fund of the Royal Society of New Zealand

## References

Baccalá, L., & Sameshima, K. (2001). Partial directed coherence: a new concept in neural structure determination. Biological Cybernetics, 84(6), 463–474.

Beier, K. T., Steinberg, E. E., DeLoach, K. E., Xie, S., Miyamichi, K., Schwarz, L., … Liqun, L. (2015). Circuit architecture of VTA dopamine neurons revealed by systematic input-output mapping. Cell, 162, 622–634.

Blanchard, T. C., Strait, C. E., & Hayden, B. Y. (2015). Ramping ensemble activity in dorsal anterior cingulate neurons during persistent commitment to a decision. Journal of Neurophysiology, 114(4), 2439–2449.

Boykin, E. R., Khargonekar, P. P., Carney, P. R., Ogle, W. O., & Talathi, S. S. (2012). Detecting effective connectivity in networks of coupled neuronal oscillators. Journal of Computational Neuroscience, 32(3), tr521-538.

Bryden, D. W., Johnson, E. E., Tobias, S. C., Kashtelyan, V., & Roesch, M. (2011). Attention for learning signals in the anterior cingulate cortex. The Journal of Neuroscience, 31(50), 18266–18274.

Carr, D. B., & Sesack, S. R. (2000). Projections from the Rat Prefrontal Cortex to the Ventral Tegmental Area: Target Specificity in the Synaptic Associations with Mesoaccumbens and Mesocortical Neurons. Journal of Neuroscience, 20(10).

Carver, C. S. (2009). Threat Sensitivity, Incentive Sensitivity, and the Experience of Relief. Journal of Personality, 77(1), 125–138. https://doi.org/10.1111/j.1467-6494.2008.00540.x

Cavanagh, J. F., Frank, M. J., Klein, T. J., & Allen, J. J. B. (2010). Frontal Theta Links Prediction Errors to Behavioral Adaptation in Reinforcement Learning. Neuroimage, 49(4). https://doi.org/10.1016/j.neuroimage.2009.11.080

Cowen, S. L., Davis, G. A., & Nitz, D. A. (2012). Anterior cingulate neurons in the rat map anticipated effort and reward to their associated action sequences. Journal of Neurophysiology, 107(9), 2393–2407.

Darwin, C. (1865). On the orgin of species by means of natural selection, or the preservation of favoured races in the struggle for life. New York: D. Appleton and Company. Retrieved from http://darwin-online.org.uk/converted/pdf/1861_OriginNY_F382.pdf

Deutsch, R., Smith, K. J. M., Kordts-Freudinger, R., & Reichardt, R. (2015). How absent negativity relates to affect and motivation: an integrative relief model. Frontiers in Psychology, 6, 152. https://doi.org/10.3389/fpsyg.2015.00152

Eshel, N., Bukwich, M., Rao, V., Hemmelder, V., Tian, J., & Uchida, N. (2015). Arithmetic and local circuitry underlying dopamine prediction errors. Nature, 525, 243–246.

Eshel, Tian, J., Bukwich, M., & Uchida, N. (2016). Dopamine neurons share common response function for reward prediction error. Nature Neuroscience, 19(3).

Faget, L., Osakada, F., Duan, J., Ressler, R., Johnson, A. B., Proudfoot, J. A., … Hnasko, T. S. (2016). Afferent Inputs to Neurotransmitter-Defined Cell Types in the Ventral Tegmental Area. Cell Reports, 15(12), 2796–2808. https://doi.org/10.1016/j.celrep.2016.05.057

Ferenczi, E., Zalocusky, K., Liston, C., Grosenick, L., Warden, M., Amatya, D., … Deisseroth, K. (2016). Prefrontal cortical regulation of brainwide circuit dynamics and reward-related behavior. Science, 351(6268), aac9698-1-aac9698-12.

Friedman, A., Homma, D., Gibb, L. G., Amemori, K., Rubin, S. J., Hood, A. S., … Graybiel, A. M. (2015). A corticostriatal path targeting striosomes controls decision-making under conflict. Cell, 161(6), 1320–1333.

Fujisawa, S., & Buzsaki, G. (2011). A 4Hz oscillation adaptively synchronizes prefrontal, VTA, and hippocampal activities. Neuron, 72, 153–165.

Gao Liu C-L., Yang, S., Jin G-Z., Bunney, B. S., Shi W-X., M. (2007). Functional coupling between prefrontal cortex and dopamine neurons in the ventral tegmental area. The Journal of Neuroscience, 27(20), 2414–5421.

Gariano, R. F., & Groves, P. M. (1988). Burst firing induced in midbrain dopamine neurons by stimulation of the medial prefrontal and anterior cingulate cortices. Brain Research, 462(1), 194–8. Retrieved from http://www.ncbi.nlm.nih.gov/pubmed/3179734

Hamid, A., Pettibone, J., Mabrouk, O., Hetrick, V., Schmidt, R., Vander Weele, C., … Berke, J. D. (2016). Mesolimbic dopamine signals the value of work. Nature Neuroscience, 19, 117–126.

Hayden, B., Pearson, J., & Platt, M. (2011). Neuronal basis of sequential foraging decisions in a patchy environment. Nature Neuroscience, 14(7), 933–939.

Hayden, B. Y., Heilbronner, S. R., Pearson, J. M., & Platt, M. L. (2011). Surprise signals in anterior cingulate cortex: neuronal encoding of unsigned reward prediction errors driving adjustment in behavior. The Journal of Neuroscience : The Official Journal of the Society for Neuroscience, 31(11), 4178–87. https://doi.org/10.1523/JNEUROSCI.4652-10.2011

Hillman, K., & Bilkey, D. K. (2010). Neurons in the rat anterior cingulate cortex dynamically encode cost-benefit in a spatial decision-making task. J Neurosci, 30(22), 7705–7713. https://doi.org/10.1523/JNEUROSCI.1273-10.2010

Howe, M., Tierney, P., Sandberg, S., Phillips, P. E. M., & Graybiel, A. M. (2013). Prolonged dopamine signalling in striatrum signals proximity and value of distant rewards. Nature, 500, 575–579.

Ito, H. T., Zhang, S.-J., Witter, M. P., Moser, E. I., & Moser, M.-B. (2015). A prefrontal-thalamo-hippocampal circuit for goal-directed spatial navigation. Nature, 522, 50–55.

Jo, Y. S., Lee, J., & Mizumori, S. J. Y. (2013). Effects of Prefrontal Cortical Inactivation on Neural Activity in the Ventral Tegmental Area. Journal of Neuroscience, 33(19).

Karalis, N., Dejean, C., Chaudun, F., Khoder, S., Rozeske, R. R., Wurtz, H., … Herry, C. (2016). 4-Hz oscillations synchronize prefrontal–amygdala circuits during fear behavior. Nature Neuroscience, 19(4), 605–612. https://doi.org/10.1038/nn.4251

Kennerly, S. W., Behrens, T. E. J., & Wallis, J. D. (2011). Double dissociation of value computations in orbitofrontal and anterior cingulate neurons. Nature Neuroscience, 14(12), 1581–1589.

Kolling, N., Behrens, T., Wittmann, M., & Rushworth, M. (2016). Multiple signals in anterior cingulate cortex. Current Opinion in Neurobiology, 37, 36–43. https://doi.org/10.1016/j.conb.2015.12.007

Likhtik, E., Stujenske, J. M., Topiwala, M. A., Harris, A. Z., & Gordon, J. A. (2014). Prefrontal entrainment of amygdala activity signals safety in learned fear and innate anxiety. Nature Neuroscience, 17, 106–113.

Ma, L., Hyman, J. M., Phillips, A. G., & Seamans, J. K. (2014). Tracking progress toward a goal in corticostriatal ensembles. Journal of Neuroscience, 34(6), 2244–2253.

Mitra, P., & Bokil, H. (2008). Observed Brain Dynamics. New York: Oxford University Press.

Morales, M., & Margolis, E. B. (2017). Ventral tegmental area: cellular heterogeneity, connectivity and behaviour. Nature Reviews Neuroscience, 18, 73–85. https://doi.org/10.1038/nrn.2016.165

Moretti, R., & Signori, R. (2016). Neural Correlates for Apathy: Frontal-Prefrontal and Parietal Cortical- Subcortical Circuits. Frontiers in Aging Neuroscience, 8, 289. https://doi.org/10.3389/fnagi.2016.00289

Navailles, S., Guillem, K., Vouillac-Mendoza, C., & Ahmed, S. H. (2015). Coordinated Recruitment of Cortical–Subcortical Circuits and Ascending Dopamine and Serotonin Neurons During Inhibitory Control of Cocaine Seeking in Rats. Cerebral Cortex, 25(9), 3167–3181. https://doi.org/10.1093/cercor/bhu112

Navratilova, E., & Porreca, F. (2014). Reward and motivation in pain and pain relief. Nature Neuroscience, 17(10), 1304–1312. https://doi.org/10.1038/nn.3811

O’Reilly, J. X., Schuffelgen, U., Cuell, S. F., Behrens, T. E. J., Mars, R. B., & Rushworth, M. F. S. (2013). Dissociable effects of surprise and model update in parietal and anterior cingulate cortex. Proceedings of the National Academy of Sciences, 110(38), E3660–E3669. https://doi.org/10.1073/pnas.1305373110

Oleson, E. B., Gentry, R. N., Chioma, V. C., & Cheer, J. F. (2012). Subsecond Dopamine Release in the Nucleus Accumbens Predicts Conditioned Punishment and Its Successful Avoidance. Journal of Neuroscience, 32(42), 14804–14808. https://doi.org/10.1523/JNEUROSCI.3087-12.2012

Onoda, K., & Yamaguchi, S. (2015). Dissociative contributions of the anterior cingulate cortex to apathy and depression: Topological evidence from resting-state functional MRI. Neuropsychologia, 77, 10–18. https://doi.org/10.1016/j.neuropsychologia.2015.07.030

Parvizi, J., Rangarajan, V., Shirer, W., Desai, N., & Greicius, M. D. (2013). The will to persevere induced by electrical stimulation of the human cingulate gyrus. Neuron, 80(6). https://doi.org/doi:10.1016/j.neuron.2013.10.057.

Paxinos, G., & Watson, C. (1997). The rat brain in stereotaxic coordinates 4th edition. New York: Academic Press.

Pearce, J. M., & Hall, G. (1980). A Model for Pavlovian Learning: Variations in the Effectiveness of Conditioned But Not of Unconditioned Stimuli. Psychological Review, 87(6), 532–552.

Qi, J., Zhang, S., Wang, H.-L., Barker, J., Miranda-Barrientos, M., & Morales, M. (2016). VTA glutamatergic inputs to nucleus accumbens drive aversion to acting on GABAeric interneurons. Nature Neuroscience, 79, 725–733.

Rajasethupathy, P., Sankara, S., Marshel, J. H., Kim, C. K., Ferenczi, E., Lee, S. L., … Deisseroth, K. (2015). Projections from neocortex mediate top-down control of memory retrieval. Nature, 526(653–659).

Rudebeck, P. H., Walton, M. E., Smyth, A. N., Bannerman, D. M., & Rushworth, M. F. (2009). Separate neural pathways process different decision costs. Nature Neuroscience, 9(9), 1161–1168.

Salamone, J. D., & Correa, M. (2012). The Mysterious Motivational Functions of Mesolimbic Dopamine. Neuron, 76(3), 470–485. https://doi.org/10.1016/j.neuron.2012.10.021

Schultz, W., Dayan, P., & Montague, P. R. (1997). A Neural Substrate of Prediction and Reward. Science, 275(5306).

Skinner, B. F. (1938). The behavior of organisms: an experimental analysis. New York: Appleton-Century-Crofts.

Sweeny, K., & Vohs, K. D. (2012). On Near Misses and Completed Tasks: The Nature of Relief. Psychological Science, 23(5), 464–468. https://doi.org/10.1177/0956797611434590

Taber, M. T., Das, S., & Fibiger, H. C. (1995). Cortical regulation of subcortical dopamine release: mediation via the ventral tegmental area. Journal of Neurochemistry, 65(3), 1407–1410.

Vogt, B. A. (2005). Pain and emotion interactions in subregions of the cingulate gyrus. Nature Reviews Neuroscience, 6(7), 533–544. https://doi.org/10.1038/nrn1704

Vogt, B. A., & Paxinos, G. (2014). Cytoarchitecture of mouse and rat cingulate cortex with human homologies. Brain Structure and Function, 219(1), 185–192. https://doi.org/10.1007/s00429-012-0493-3

Walton, M. E., Devlin, J. T., & Rushworth, M. F. (2003). Functional specialization within medial frontal cortex of the anterior cingulate for evaluating effort-related decisions. Journal of Neuroscience, 23, 6475–6479.

Wu, J., Gao, M., Shen, J. X., Shi, W. X., Oster, A. M., & Gutkin, B. S. (2013). Cortical control of VTA function and influence on nicotine reward. Biochem Pharmacol, 86(8), 1173–1180. https://doi.org/10.1016/j.bcp.2013.07.013

